# Dynamic encoding of valence by anterior cingulate cortex in mice

**DOI:** 10.1101/2025.06.17.660226

**Authors:** Priyodarshan Goswamee, Nilanjana Saferin, Radha Shah, Jorge S. Seoane, Kamalika Ganguly, Kari L. Neifer, Robert A. Pearce, James P. Burkett

**Author notes:** Corresponding authors: James P. Burkett, Ph.D. 3000 Arlington Ave, Block Health Sciences #185 Toledo, OH 43614, Priyodarshan Goswamee, Ph.D. 1101 East Marshall St., Sanger Hall, Room 12-067A Richmond, VA 23298.

## Abstract

Fear supports survival by linking environmental cues with potential threats, but its dysregulation contributes to anxiety disorders and PTSD. While the contributions of hippocampus and amygdala to fear memory are well established, the role of the anterior cingulate cortex (ACC) during and after acquisition remains less clear. Here we longitudinally recorded ACC neuronal activity using single-photon calcium imaging in freely behaving mice undergoing auditory fear conditioning. During cue pre-exposure, neural responses to the cue were strongest to early novel presentations and declined with repetition. During acquisition, responses emerged late in training as the cue became shock-predictive. Ensemble analysis identified subpopulations of “freezing on” and “freezing off” cells in the ACC whose activation coincided with decreased neural activity and increased neural synchrony. During 24-h and 48-h recall, ACC responses were robust to early presentations but diminished during late tones; similarly, a subpopulation of neurons emerged that was upmodulated by the CS in early 24-h and 48-h recall. Together, our results demonstrate that ACC neuronal response to a cue reorganizes both within and across sessions, with dynamic changes in population activity, recruitment, and synchrony that mirror changes in novelty and negative valence. This pattern suggests a role for ACC as a flexible hub that dynamically represents changes in cue significance.

## Introduction

Fear memory enables learned associations between environmental cues and potential threats, and its dysregulation can lead to debilitating conditions such as anxiety disorders and post-traumatic stress disorder (PTSD) (Bowers & Ressler, 2015). Understanding the neurobiological substrates of fear memory is crucial for addressing these conditions. In mammals, fear memory is neuronally represented by the “fear engram”, which resides in the basolateral amygdala and hippocampus. The anterior cingulate cortex (ACC), a part of the prefrontal cortex with strong synaptic connections to amygdala and hippocampus, has emerged as a key node in aversive learning (Feng et al., 2014; Pissiota et al., 2003; Toyoda et al., 2011; Bissière et al., 2008; de Lima et al., 2022; Goshen et al., 2011; Han et al., 2003; Jhang et al., 2018; Kitamura et al., 2017; Steenland et al., 2012; Tang et al., 2005). Direct stimulation of the ACC in mice induces freezing responses and consolidates long-term fear memory, while transient inactivation impairs associative fear learning, underscoring its role in the acquisition of fear (Bissière et al., 2008; de Lima et al., 2022; Tang et al., 2005). Some research suggests that the ACC is exclusively involved in recall of remote memories (~30 days) (Frankland et al., 2004; Goshen et al., 2011), while others suggest a broader role for the ACC, including the encoding and reconsolidation of recent (24-h) fear memories (Einarsson & Nader, 2012). Thus, ACC’s precise contributions across different temporal stages of fear memory remain unresolved.

In recent years, refined neurotechnologies such as optogenetics, single- and two-photon calcium imaging, have led to a surge in the exploration of the relationship between brain network activity and behavior. Single-photon calcium imaging in freely behaving rodents offers the opportunity to longitudinally track the activity of hundreds of neurons of a target brain region (Patel et al., 2022). Given evidence that ACC encodes prediction errors (Alexander & Brown, 2019) and is modulated by attention (Wu et al., 2017) and motivation (Rolls, 2019; Vassena et al., 2020), we hypothesized that the ACC supports cue discrimination by enhancing representations of cues that predict negative outcomes. Here, we performed single-photon calcium imaging in seven freely-behaving adult male mice throughout a multi-day cued fear conditioning paradigm (context pre-exposure, cue pre-exposure, fear acquisition, and memory tests at 24 and 48 hours). We measured single-cell and population activity from >2,000 longitudinally registered ACC neurons during behavioral states (movement and freezing) and exposure to a conditioned stimulus (CS, tone) and an unconditioned stimulus (US, light foot shock), across all stages of the experiment. In brief, we found that ACC neurons showed dynamic modulation by both the conditioned and unconditioned stimuli independent of behavior-related activity changes, with patterns suggesting ACC encodes the changing significance of the cue. Our work offers new insights into the role of the ACC in fear memory processing and may help identify potential cortical targets for interventions.

## Methods

### Subjects

Experimental subjects were adult C57BL/6J mice, 4-6 months of age, bred in our animal facility, and housed in a temperature- and humidity-controlled housing facility under a 12-h light/dark cycle with *ad libitum* access to food and water. Following surgery, animals were singly housed to prevent potential injury to their implant site. All experiments were approved by the Institutional Animal Care and Use Committee of the University of Toledo and were conducted as per the recommendations of the National Institute of Health guidelines for the Care and Use of Laboratory Animals.

The a priori elimination criteria were (1) implant placement outside the ACC and (2) fewer than 50 longitudinally registered neurons. Male (N=8) and female (N=3) mice underwent implant surgery as described below. One male subject was eliminated for having too few longitudinally registered neurons, and all three females ended up being excluded due to poor implant placement, yielding a final cohort of N=7 male mice.

### Surgery

To conduct single-photon calcium imaging, we intracranially injected adeno-associated viruses (AAV) that expressed GCaMP6f in excitatory cells (AAV1.CamK2a.GCaMP6f.WPRE.bGHpA; Inscopix, Mountain View, CA) into the right ACC and implanted a 1-mm (*dia.*) X 4-mm GRIN lens (Proview Integrated GRIN lens, Inscopix) over the GCaMP6f-expressing neurons in the ACC, enabling transmission of the fluorescent signal to the miniscope (nVoke 2.0, Inscopix) docked onto a magnetic base plate cemented to the animal’s skull. For stereotactic AAV injection and implantation of GRIN lenses, we anesthetized (100 mg/kg ketamine and 2.5 mg/kg xylazine intraperitoneal [IP]) and mounted mice on a stereotactic frame equipped with a robotic controller (NeuroStar, Sindelfingen, Germany). We maintained anesthesia with 1-2% oxygen-vaporized isoflurane (isoflurane: Covetrus, Portland, ME; isoflurane vaporizer: R583S, RWD, Shenzhen, China; oxygen concentrator: ROC-5A, RWD, Shenzhen, China) and monitored body temperature (Thermostar 69020, RWD, Shenzhen, China) for the duration of the surgery. All animals received subcutaneous injections of atropine (0.04 mg/kg) and extended-release analgesia (Ethiqa XR, 0.05 ml) at the start of the surgery to control excessive secretion and postoperative pain, respectively. We shaved and prepped the incision site with an alcohol and iodine scrub, exposed the skull via a midline incision, and treated with topical lidocaine. A small hole (1.8 mm diameter) was drilled in the skull overlying the right ACC (AP +1.0 mm, ML +0.4 mm) using a trephine bur attached to a motorized drill. Following excision of the dura, the cortex was aspirated up to a depth of 0.5 mm using a 27G blunt-end needle (SAI Infusion Technology, Lake Villa, IL) accompanied by continuous perfusion with ice-cold, sterile lactate Ringer’s solution. We lowered an aluminosilicate glass pipette containing the virus to the level of the right ACC at DV −1.75, −1.80, and −1.85 mm and infused 500 nl virus (titer strength: 3.35 × 10^12^ vg/mL) at a rate of 100 nl/min at each depth using a Neurostar-driven Injectomate, after which the injection pipette remained in place for 5 min before withdrawal. A 1-mm diameter x 4-mm long GRIN lens integrated with a baseplate (Inscopix) was lowered at 100 µm/min to the ACC above the viral injection site (AP +1 mm; ML +0.5 mm; DV 1.7 mm) and fixed in place using Metabond cement (Parkel, Edgewood, NY). A silicon adhesive (Kwik sil, World Precision Instruments, Sarasota, FL) was applied to cover the exposed section of the brain before Metabond application. All animals were administered an IP injection of 0.7-1.0 ml warm lactate Ringer’s solution before waking up. Following surgery, all animals were individually housed in draft-free home cages with *ad libitum* access to enriched nourishment (Boost diet gel and sunflower seeds) and water. Animals were allowed to recover for 4 weeks and checked for expression of GCaMP6f before the commencement of behavioral habituation.

### Single-photon Calcium Imaging

For calcium (Ca²⁺) imaging, GCaMP6f fluorescent signals were captured at 60 frames/s by cycling through three focal planes (20 Hz per plane) using 488-nm excitation light delivered though the GRIN lens by a miniscope (nVoke 2.0, Inscopix). The light beam intensity was set between 0.7 and 0.9 mW/mm². Simultaneous recording of calcium signals, animal behavior, and delivery of the stimuli were coordinated between the Inscopix Data Analysis Software (Inscopix), the nVision behavioral recording system (Inscopix), and the Freezeframe software (Actimetrics, Wilmette, IL) via Transistor-to-Transistor Logic (TTL) pulses.

### Fear Conditioning

Figure 1A provides the experimental timeline and a graphical representation of the setup. Following 4 weeks of recovery from surgery, mice habituated to the head-mounted miniscope that is tethered via an optic fiber cable to a commutator (Inscopix) for 3 consecutive days, during which the animal spent 5 min (during the first three days) or 15 min (last 2 days) inside a fear conditioning chamber (Coulbourn Instruments, Allentown, PA) in a sound-attenuating chamber (Harvard Apparatus) positioned inside a second sound-attenuating cabinet (ROOM, New York, NY). Behavior was recorded using the nVision camera (Inscopix).

**Figure 1.**
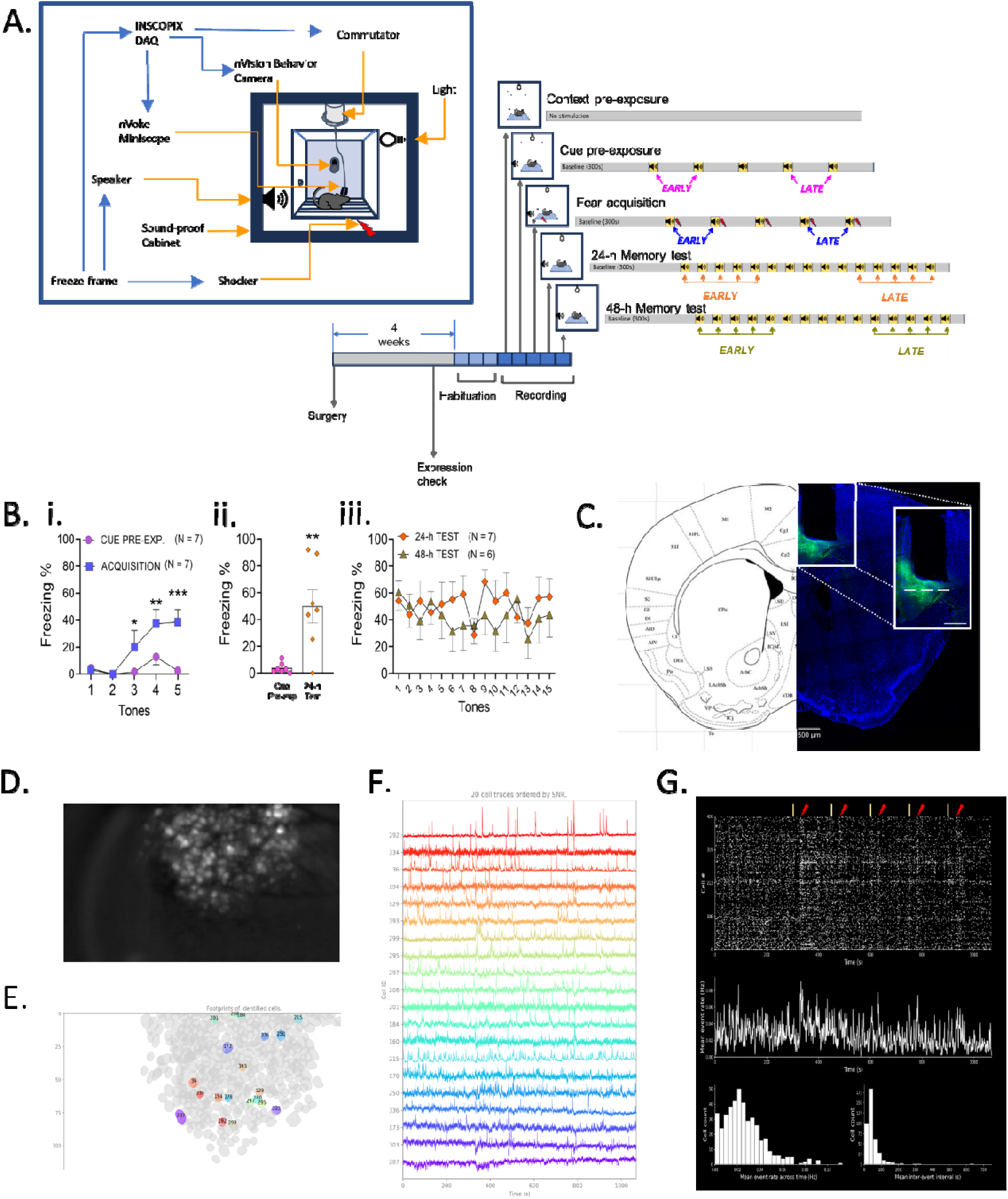
Experimental details of miniscope imaging in the mouse ACC. **A:** Schematic of the experimental setup (left) and study timeline (right). Blue arrows represent TTL signal routing for synchronized acquisition. The behavioral paradigm included context pre-exposure, cue pre-exposure, fear acquisition, and subsequent 24-h and 48-h memory tests. The “Early” and “Late” tones within each session are indicated. **B:** Within-subjects freezing in response to tone cues presented during cue pre-exposure, acquisition, and 24-h and 48-h memory sessions. **i.** Acquisition of the conditioned freezing response. **ii.** Recall of the conditioned response in the first 5 tones of the 24-h memory test. **iii.** Lack of behavioral extinction across 30 tones in 24-h and 48-h tests. Error bars are SEM. * p<0.05; ** p<0.01; *** p<0.001. **C:** A representative coronal atlas image (left) and fluorescent section (right) showing viral expression of GCaMP6f and GRIN lens placement in the right ACC. The magnified white inset highlights the imaging plane (dashed line). Scale bar = 0.5 mm (inset = 1 mm). **D:** Maximum projection of a calcium imaging movie from a representative subject, showing all active neurons in a single session. **E:** Cell map from the same subject, generated using CNMF-E, showing spatial footprints of identified neurons. **F:** ΔF/F traces from 20 neurons ordered by signal-to-noise ratio (SNR); colors correspond to ROIs in panel E. **G:** Top: Raster plot of calcium events recorded during the acquisition session (day 3). Yellow and red ticks mark CS (tone) and US (shock) onset, respectively. Middle: mean population event rate across the session. Bottom: histograms showing distribution of mean event rates (left) and inter-event intervals (right).

The following week, we recorded behavior concurrently with neuronal activity in the ACC as the mice underwent a 5-day fear conditioning protocol. On day 1 (context pre-exposure), the mice freely explored the conditioning chamber for 15 min. On day 2 (cue pre-exposure), following 5 min of habituation, subjects were exposed to five presentations of the conditioned stimulus (CS), which consisted of a neutral tone cue (30 s, 6000 kHz, 85 dB, 2-min inter-trial interval). On day 3 (fear acquisition), subjects experienced the same protocol as day 2 except that the unconditioned stimulus (US), consisting of a mild foot shock (0.8 mA, 0.5 s), was delivered 29.5 s after the onset of each CS. On day 4 (24-h test) and day 5 (48-h test), subjects were placed in the conditioning chamber and then, after 5 min of habituation, exposed to 15 presentations of the CS at 30-s inter-trial intervals. A within-session movement control period (first 300 s) was included at the beginning of every session to measure baseline activity in a neutral state.

Conditioned freezing was assessed by quantifying mouse freezing behavior following CS onset on days 2 through 5. Behavioral analysis was performed offline using Ethovision XT 16 (Noldus), which created an annotated behavior trace with two defined states: “Freezing” and “Moving.” Freezing bouts were defined operationally as movement below a threshold of 0.01% for longer than 1 s. Technical issues prevented data collection for one subject during the day 5 session. Additionally, only a partial dataset (limited to the first three tones) was available from the day 2 session for another subject. The percentage of freezing during CS presentations was compared among all subjects using mixed-effect models (MEM) due to availability of partial datasets for some of the recording sessions. An omnibus MEM with repeated measures of day and tone was used, followed by post-hoc MEMs comparing cue pre-exposure to acquisition (acquisition), pre-exposure to the first 5 tones of the 24-hour test (recall), and 24-hour test to 48-hour test (extinction). Additional post-hoc Tukey’s tests compared tones between days.

### Verification of virus expression and lens placement

At the end of behavioral experiments, animals were euthanized by isoflurane overdose followed by exsanguination via transcardial perfusion. Perfusion was performed using PBS followed by ice-cold 4% paraformaldehyde (PFA) solution. The brains were removed and submerged into 4% PFA for another 24-h at 4°C. Fixed brains were then transferred into a 30% sucrose solution. After sinking to the bottom of the storage container, the brains were embedded into plastic molds with OCT gel, rapidly frozen using dry ice, and stored at −80°C. The frozen tissue blocks were later sectioned using a cryostat (CM3050S, Leica Microsystems, Wetzlar, Germany). Coronal sections (40-µm) containing the ACC were mounted on Superfrost+ glass slides using Vectashield mounting medium and imaged using a fluorescent microscope with 4X magnification. Imaged sections were overlaid on top of an online mouse brain coronal atlas (http://labs.gaidi.ca/mouse-brain-atlas/) to confirm the position of the implanted lens and GCaMP6f expression.

### Primary Processing of the Calcium Imaging Dataset

The miniscope movies were initially processed using the Inscopix data processing system (IDPS Version 1.8). The Ca^2+^ imaging movie files from three focal planes were spatially and temporally down-sampled by a factor of two to generate movie files (320 *X* 198 pixels, 10 Hz frame rate). The movies were further processed using a Gaussian filter and motion-corrected for neuron identification. The Constrained Nonnegative Matrix Factorization for microendoscopic data (CNMFe) algorithm (Zhou et al., 2018) was used to extract the spatial footprints of the neurons. For CNMFe, when searching for new cell seeds, we set we set the minimum seed criteria as: 0.8 Pearson correlation with neighbors; Signal-to-noise ratio (SNR) ≥ 10; and ~10 μm cell diameter. Multiplane registration of the coordinates of the seed pixel positions was utilized to obtain a single cell set containing every neuron from all three focal planes. The change in GCaMP6f fluorescence over time was computed and represented as ΔF/F traces, which were further analyzed using the Inscopix Data Exploration, Analysis, and Sharing (IDEAS) Platform. Positive deflections in the ΔF/F traces that showed a fast increase in amplitude followed by a long decay back to baseline level were identified by the software as putative Ca^2+^ events. The default value of the event’s smallest decay time was set at 0.2 s in the algorithm, as measured from the response of GCaMP6f corresponding to one action potential, as established by others (Chen et al., 2013). The threshold used to screen for events was empirically determined and set at 6x standard deviation. Raw traces were examined individually and accepted as a cell if the corresponding neurons showed activity >0.1 Hz and cell diameter between 5 and 30 µm. The Ca^2+^ events were synchronized with behavioral states (mobile or freezing) via common timestamps or TTL pulses generated by Ethovision XT software.

### Data Analysis and Statistics

#### Neural activity during behavioral states

The Compare Neural Activity Across module in IDEAS (v2.1.0) was used to compute the mean calcium event rate (population activity) during each behavioral state (‘freezing’ vs ‘moving’) using the LR cell set for acquisition vs. 24-h test. For each epoch, the module outputs mean population activity traces, individual cell activity, and categorization of cells as moving-preferring, freezing-preferring, or non-preferring. Mean population activity was compared across states and sessions using an ANOVA with post-hoc t-tests. Proportions of state-preferring cells, and the transitions between preferences across sessions, were compared using McNemar’s tests with φ coefficient. Relative neural activity of moving-preferring and freezing-preferring cells were calculated by subtracting the neural activity of non-preferring cells for each behavioral state and compared using an ANOVA with post-hoc t-tests.

#### Neural circuit correlations

The Compare Neural Circuit Correlations Across States algorithm (ver. 7.1.3) of the IDEAS package was used to evaluate pairwise correlations of activity during the first 10 s after the onset of an event or series of events (tones for cue-evoked analyses, or the start of a behavioral state episode for state analyses) using the LR cell sets. A frequency distribution of the maximum correlation for all LR cells was generated using the 10 s post-event window or the behavior state. A reference distribution (“Other”) for each session was computed from non-event-aligned neural activity windows selected to exclude stimulus-associated epochs, providing a baseline for spontaneous or background correlation levels. Frequency distribution histograms (0.05 unit bins) were computed for each subject and each epoch within each LR cell set, and the subject-level histograms were compared between states and to the reference distributions using ANOVAs. Frequency distributions were graphed as cumulative frequency distributions (CFD). The Compare Neural Circuit Correlations Across States algorithm also generated post hoc analyses for positively correlated and negatively correlated pairwise activity.

#### Peri-event analysis

We used the Peri-Event Analysis algorithm (ver. 5.3.1 & 5.6.1) and the Combine and Compare algorithm (ver. 2.5.1) in the IDEAS platform, which compute the change in neural activity within a −20 s to +20 s window before and after a single event (*e.g., a* CS or US) or a series of events, to analyze the change in firing activity of both the entire population of neurons and each individual neuron and to classify them into up-modulated, down-modulated, or non-modulated sub-populations. For each analysis, a null distribution for comparison was generated by circularly permuting the event time relative to the neural time series using 1000 random shuffles, resulting in a p-value resolution of α=0.001. The neural activity Z-score of individual neurons was calculated at each time increment using the equation Z = (*X* − µ)/a where X is the observed firing rate, μ is the mean firing rate, and σ is the standard deviation of the firing rate. Next, the bootstrap probability was calculated by determining the fractions of the null distribution that were greater than and less than the observed neural activity Z-score. Two-tailed t-tests were performed, and populations/cells were labeled as up-modulated if the bootstrap p-value was <α/2, down-modulated if the bootstrap probability was >1-α/2, and non-modulated if the bootstrap probability was between α/2 and 1-α/2 (α=0.001). LR cell sets were used to compare between sessions of interest (Supplementary Fig. S2D), while all accepted cells from a session were used for within-session comparison of activity. Peri-event analyses were conducted separately for each LR cell set and for each within-session all accepted cell set. The proportions of up-, down- and non-modulated cells, as well as the transitions between modulation states within and between sessions, were compared using McNemar’s tests with φ coefficient.

The event-associated population activity (EAPA) traces were exported and processed using a custom Python script that performed the following calculations. EAPA traces were smoothed into a 5-point moving average for noise reduction. Baseline activity was defined as the mean population activity from −20 to 0 s relative to event onset. Peak amplitude (ΔZ*_PEAK_*) was defined as the difference in Z-score between the post-event maximum (from 0 to +20 s) and baseline activity. Peak onset and peak offset were defined as the closest time points before and after the peak, respectively, at which population activity returned to baseline activity, constrained to the 0 s to +20 s event window. The response magnitude was defined as the area under the curve (AUC) of the baseline-subtracted population activity between peak onset and peak offset. Peak amplitudes and response magnitudes were compared between epochs using t-tests.

#### Statistics

All statistical analyses were performed using SPSS v30.0.0 (IBM, Armonk, NY) and Prism v10.6.0 (GraphPad Software, Boston, MA). As all measures were within-subjects, all ANOVAs and mixed models used within-subjects factors, and all post-hoc tests were paired. Similarly, all tests of cell proportions used within-subjects McNemar’s tests, with the φ coefficient for evaluating symmetric vs. asymmetric transitions between states. For evaluation of observed vs. expected proportions, the expected percentages of up-modulated, down-modulated and non-modulated neurons by chance were considered to be 2.5%, 2.5%, and 95%, respectively.

## Results

### Classical fear conditioning

We used a 5-day auditory fear conditioning paradigm consisting of context pre-exposure, cue pre-exposure, acquisition, and memory tests at 24-h and 48-h, with tone cues as the CS and light foot shocks as the US (Fig. 1A). This design allowed us to examine both the behavioral expression of fear and the longitudinal dynamics of anterior cingulate cortex (ACC) population responses across learning and recall. As expected, freezing behavior during CS presentations varied significantly across days (MEM, main effect of day, F(3,18)=15.81, p<0.0001; Fig. 1B, Fig. S1). Freezing behavior increased during acquisition relative to pre-exposure over the course of 5 CS presentations (day*time interaction, F(4,22)=4.31, p=0.010; post-hoc Tukey’s tests: CS 3, p=0.030; CS 4, p=0.0053; CS 5, p=0.0001), demonstrating successful acquisition of a conditioned freezing response. Freezing remained elevated during the first 5 CS presentations on the 24-h test day (main effect of day, F(1,6)=15.37, p=0.0078), demonstrating successful recall of the conditioned response. However, freezing did not decrease over the course of 30 CS presentations in the 24-h and 48-h tests (no main effect of time, no main effect of day, no time*day interaction), with freezing remaining consistent from the first 5 presentations in the 24-h test to the last 5 presentations in the 48-h test (F(1,6)=0.069, p=0.80), suggesting that extinction of the conditioned response did not run to completion within this behavioral paradigm.

### Viral expression, lens placement, and registration of neurons

Prior to behavioral testing, mice underwent stereotaxic surgery for the implantation of a GRIN lens and injection of a GCaMP6f-expressing viral vector targeting the right ACC. Histological analysis confirmed robust virus expression and correct lens placement in all included animals (Fig. 1C), yielding a final cohort of N=7 mice. We extracted spatial footprints of ACC neurons and performed multi-plane registration to track cells across the three imaging planes (Fig. 1D-E; Fig. S2A-D). Data were processed and calcium events detected using the IDEAS platform. Raw calcium traces, annotated raster plots, and event timing distributions from a representative subject during the fear conditioning session are shown in Figs. 1F and 1G. Within-session comparisons used the full multi-plane registered cell sets for each session, while between-sessions comparisons used only longitudinally registered cells between the sessions being compared (Fig. S2D).

### ACC subpopulations represent moving and freezing states

We first investigated whether the activity of ACC neurons represented the behavioral state. The calcium activity traces during the acquisition and 24-h test sessions, representing neural activity, were overlaid with behavioral traces with freezing and moving states annotated (Figs. 2A-B). Neural activity in ACC across both sessions depended on the behavioral state (ANOVA, main effect of state, F(1,6)=9.58, p=0.021), with a relative decrease in neural activity in the freezing state (t-test, p=0.021; Fig. 2C). Individual neurons were then categorized as “moving-preferring,” “freezing-preferring,” or “non-preferring” based on their neural activity during both states. The observed proportions of moving-preferring and freezing-preferring neurons in both sessions were greater than expected by chance (McNemar’s test: acquisition, χ²=3081, p<0.0001; 24-h test, χ²=1695, p<0.0001), although the proportion of moving-preferring neurons diminished in the 24-h test session (McNemar’s test, χ²=26.23, p<0.001; Fig. 2D). Neurons transitioned between firing preferences asymmetrically between sessions (φ=0.061, p=0.037), with both moving-preferring and freezing-preferring neurons maintaining their firing preferences across sessions (Fig. 2E). Neural activity in the two stable subpopulations differed depending on the behavioral state (ANOVA, state*preference interaction, F(2,12)=23.38, p<0.001), with neural activity in moving-preferring neurons being suppressed during the freezing state (t-test, p=0.023) and neural activity in freezing-preferring neurons being elevated during the freezing state (t-test, p=0.0034) (Fig. 2F). Therefore, these cell populations could be referred to as “Freezing Off” and “Freezing On,” respectively.

**Figure 2.**
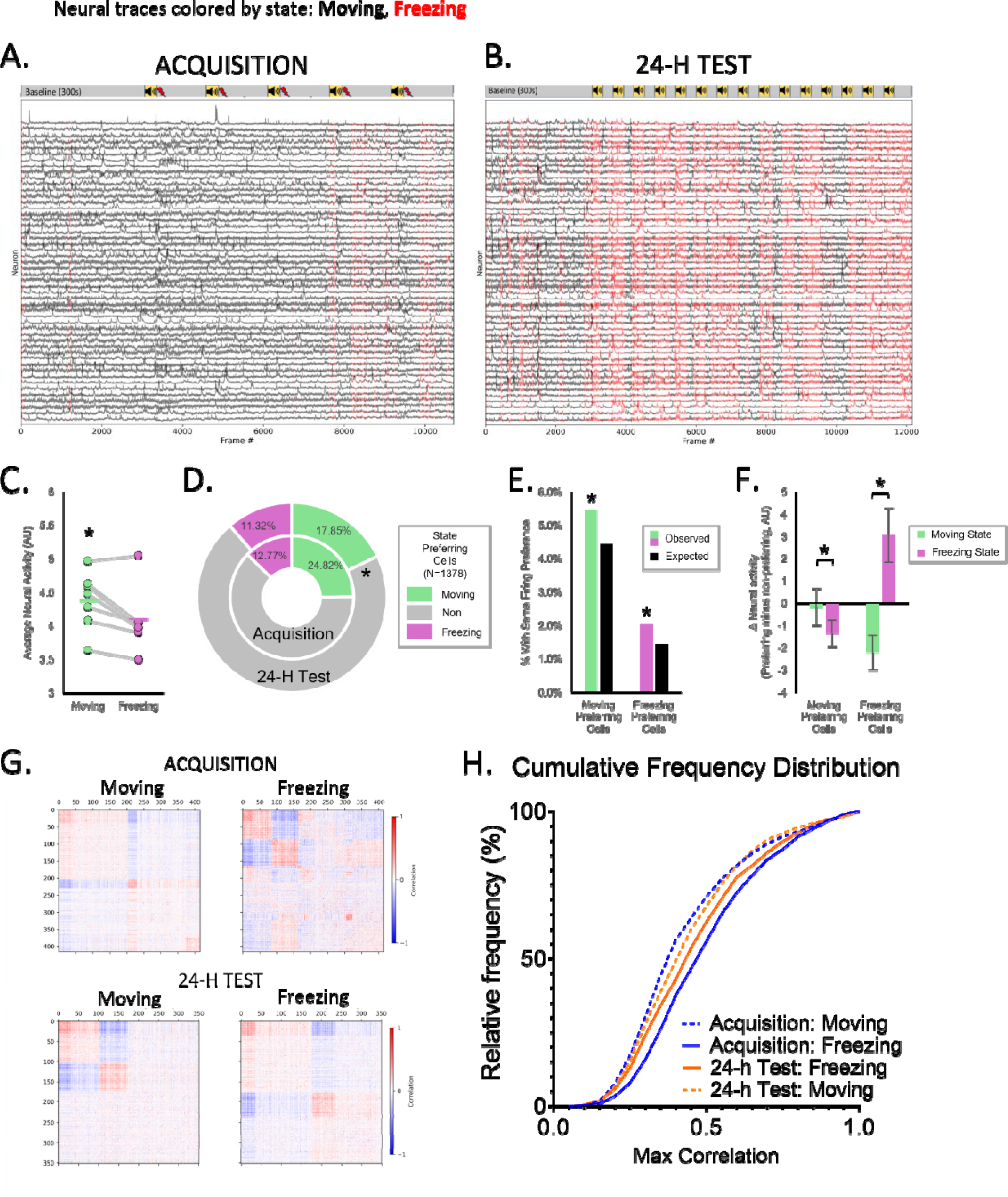
Neural response comparison between moving and freezing states. **A:** Representative calcium traces from a single mouse during the acquisition session, with traces colored according to behavioral state (black = moving, red = freezing). **B:** Representative calcium traces from the 24-h test session. **C:** Mean neural activity of LR cells from both sessions separated by state. **D:** The proportions of up-, down-, and non-modulated neurons during moving and freezing states in the acquisition and 24-h test sessions. **E:** Observed vs. expected proportions of neurons maintaining their firing preferences between sessions. **F:** Neural activity of moving-preferring and freezing-preferring neurons relative to non-preferring neurons in each behavioral state. **G:** Correlation heatmaps of all neurons during moving and freezing in the acquisition and 24-h test sessions. **H:** Cumulative frequency distributions of maximum pairwise correlation values in moving and freezing states in both sessions. Error bars are SEM. * p<0.05.

To address whether neural activity in the ACC consistently predicted (and therefore potentially drove) the onset of behavioral freezing, we compared peri-event activity for freezing onsets in the no-tone period (context alone) vs. freezing onsets during CS presentations (Fig. S3A). Averaged population activity aligned to freezing onset did not differ between groups (Up-modulated: p = 0.39; Down-modulated: p = 0.66; Fig. S3B). Likewise, the proportions of up-, down-, and non-modulated neurons were statistically indistinguishable (χ² = 0.72, p = 0.70; Fig. S3C). These findings suggest that ACC neural responses are not sufficient to predict freezing onsets, and therefore that ACC is unlikely to directly drive freezing behavior.

To assess ensemble dynamics, we calculated the frequency distribution of pairwise correlated neuronal activity (Figs. 2G-H). Correlated neuronal activity differed across behavioral states, with stronger correlations being more frequent in the freezing state (ANOVA, state*correlation interaction, F(18,108)=2.28, p=0.005; Fig. 2H). This rightward shift in correlated activity during the freezing state was greater during acquisition (day*state*correlation interaction, F(18,108)=3.35, p<0.001; paired t-tests, p<0.05) and was not specific to positive or negative correlations (all p>0.6; Fig. S4). These results show an increase in neural synchrony in the freezing state, particularly during acquisition.

Overall, these results show stable subpopulations of “Freezing Off” and “Freezing On” cells in the ACC during acquisition and 24-h test sessions, together with decreased neural activity and increased neural synchrony during the freezing state.

### ACC represents aspects of the US

We next analyzed how ACC neurons represented the US during fear acquisition, consistent with its known role in nociception and aversion processing (Niikura et al., 2011; Zhao et al., 2018; Acuña et al., 2023; Li et al., 2023). The US consistently evoked a response in the event-aligned population activity (EAPA) (Figs. 3A-B; Fig. S5), and the proportions of both up-modulated and down-modulated neurons across all US events were higher than expected by chance (χ^2^=263; p<0.00001; Fig. 3C). The EAPA peak amplitude differed significantly across US events (ANOVA, F(2.3,13.9)=5.2, p=0.017, Fig. 3D), with the highest amplitude response to US 1. The pattern of change in the response magnitude, calculated by integrating the area under the curve (AUC) between peak onset and peak offset, was superficially similar but did not differ significantly across US events (ANOVA, p=0.3196, F(2.3,13.7)=1.3; Fig. 3E). The proportions of modulated neurons also differed across US events (McNemar’s test, X2=127.8, p<0.001), with US 1 recruiting significantly more up-modulated neurons than the other US events, and US 3 recruiting significantly fewer up-modulated and down-modulated neurons (Fig. 3F). The response amplitudes did not differ significantly across US events for either up-modulated or down-modulated groups (all p>0.5; Fig. S6), suggesting that the primary driver of differences in response amplitude across US events was the number of modulated neurons. These results are consistent with a consistent representation of the aversive US, concurrent with an initial, rapidly attenuating representation of US novelty/prediction error.

**Figure 3.**
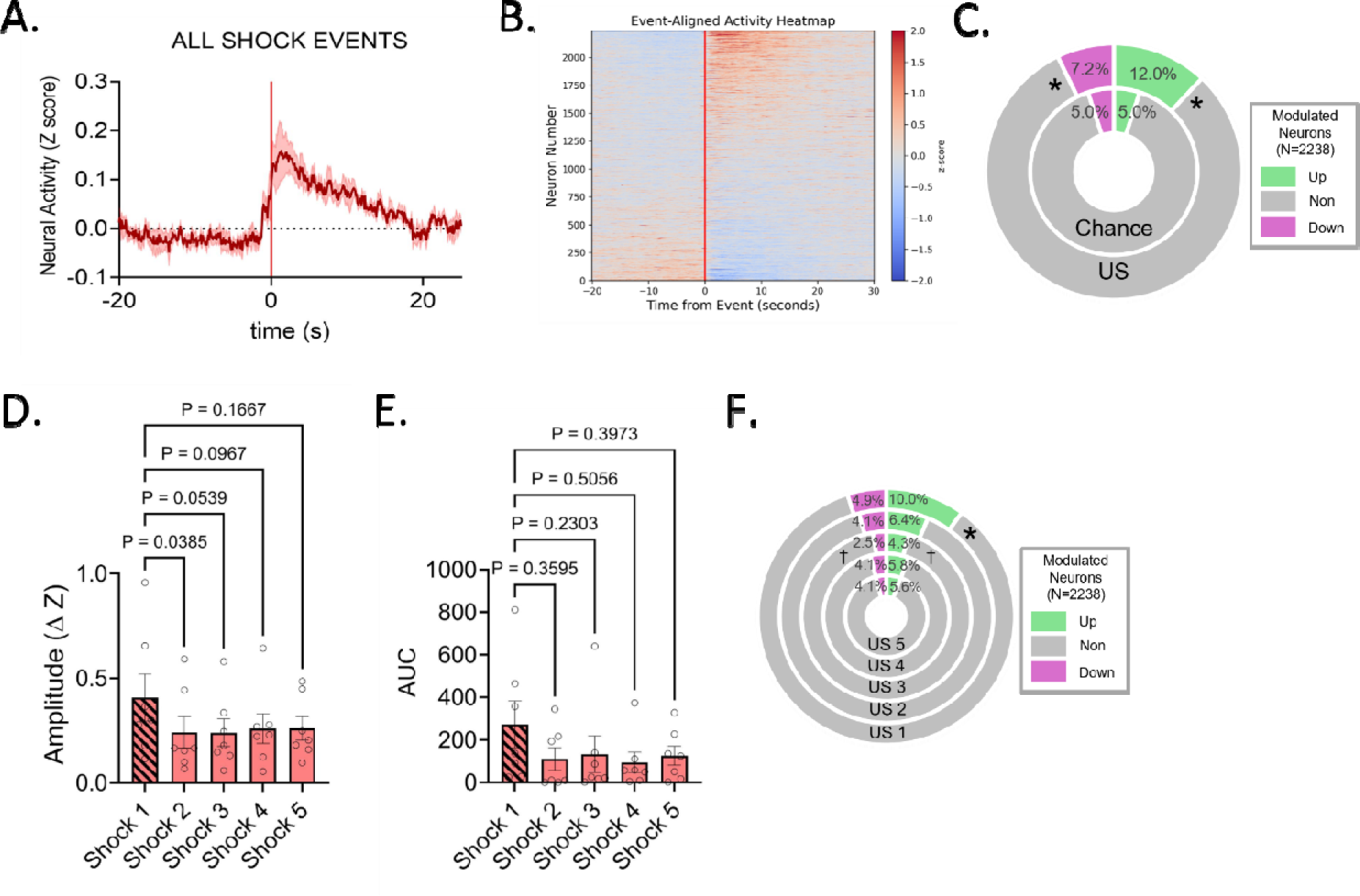
Neural activity during US events. **A:** Event-aligned population activity trace (EAPA) showing mean ACC activity across all US events. **B:** Heatmap of trial-by-trial activity aligned to US onset, ordered by maximum response. **C:** The proportions of up-, down-, and non-modulated neurons during all US events combined, as compared to the proportions expected by chance. * p<0.05. **D-E:** Quantification of US responses across the five individual US events, showing (D) peak amplitude and (E) response magnitude (AUC). **F:** The proportions of up-, down-, and non-modulated neurons during each of the five individual US events. All error bars are SEM. * higher proportion, p<0.05; † lower proportion, p<0.05.

### ACC dynamically represents multiple characteristics of the CS

We next examined how ACC responses to the CS evolved across cue pre-exposure, acquisition, and the 24-h and 48-h memory tests (Figs. 4, 5, S7). Within each cue session, tones were classified as “early” and “late” to capture potential differences in ACC activity as cue significance evolved across trials. Comparing responses between cue pre-exposure and acquisition sessions by averaging across all five tones showed no significant differences in peak amplitude, AUC, or fraction of modulated neurons (all p>0.1; Fig. S7). In contrast, separating early and late tone responses revealed strong, dynamic changes in neural responses both within and between sessions. The first presentations of the CS during early cue pre-exposure elicited strong EAPA responses that rapidly attenuated, with amplitude, magnitude, and proportions of modulated neurons all declining from early to late tones (all cells; amplitude: t-test, p=0.043; magnitude: t-test, p=0.043; proportions: McNemar’s test, χ^2^=22.1, p<0.001; Figs. 5A-C). EAPA responses remained low during early acquisition, with fewer neurons modulated as compared to early pre-exposure (LR cells; χ^2^=42.90, p<0.001; Figs. 4A-D). By contrast, late acquisition tones showed the opposite pattern: EAPA response magnitude and proportions of modulated neurons rose from early to late acquisition (all cells; magnitude: p=0.0156; proportions: χ^2^=13.38, p=0.004; Figs. 5A-C), and both peak amplitude and magnitude were significantly greater during late acquisition compared to late pre-exposure (LR cells; amplitude: p=0.0053; magnitude: p=0.014; Figs. 4E-H). These results suggest that ACC neurons show greater responses to early tones during cue pre-exposure, consistent with its role in novelty-related prediction error; and while novelty-related responses decline over time, responses increase in late acquisition after the cue is paired with the US.

**Figure 4.**
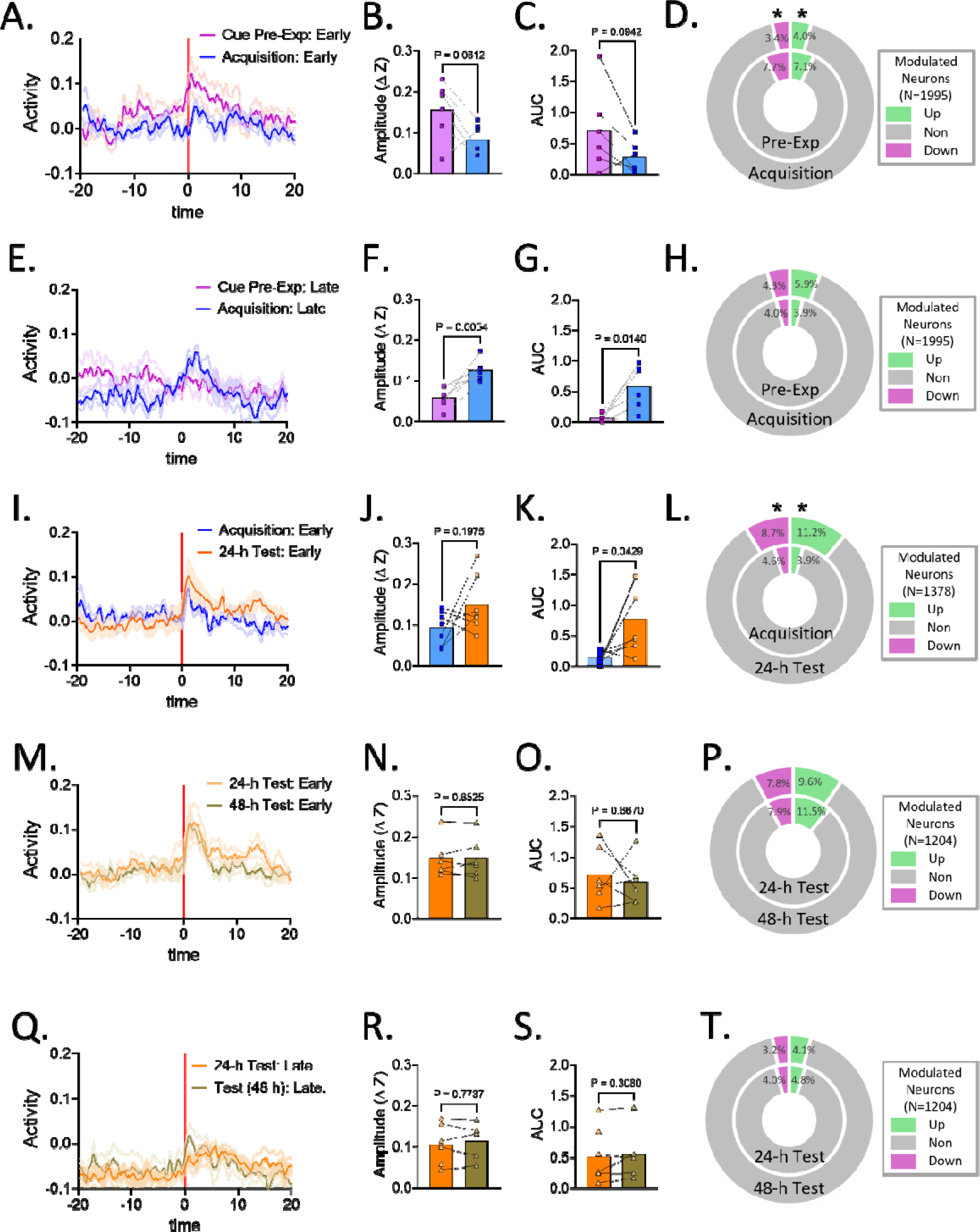
Dynamic changes in neural responses between sessions. **A-D:** Early tones during cue pre-exposure vs. acquisition compared on (A) event-aligned population activity (EAPA) traces, (B) peak amplitude, (C) response magnitude, and (D) proportions of modulated neurons. **E-H:** Late tones during cue pre-exposure vs. acquisition compared on (E) EAPA traces, (F) peak amplitude, (G) response magnitude, and (H) proportions of modulated neurons. **I-L:** Early tones during acquisition vs. 24-h test compared on (I) EAPA traces, (J) peak amplitude, (K) response magnitude, and (L) proportions of modulated neurons. **M-P:** Early tones during 24-h test vs. 48-h test compared on (M) EAPA traces, (N) peak amplitude, (O) response magnitude, and (P) proportions of modulated neurons. **Q-T:** Late tones during 24-h test vs. 48-h test compared on (Q) EAPA traces, (R) peak amplitude, (S) response magnitude, and (T) proportions of modulated neurons.

**Figure 5.**
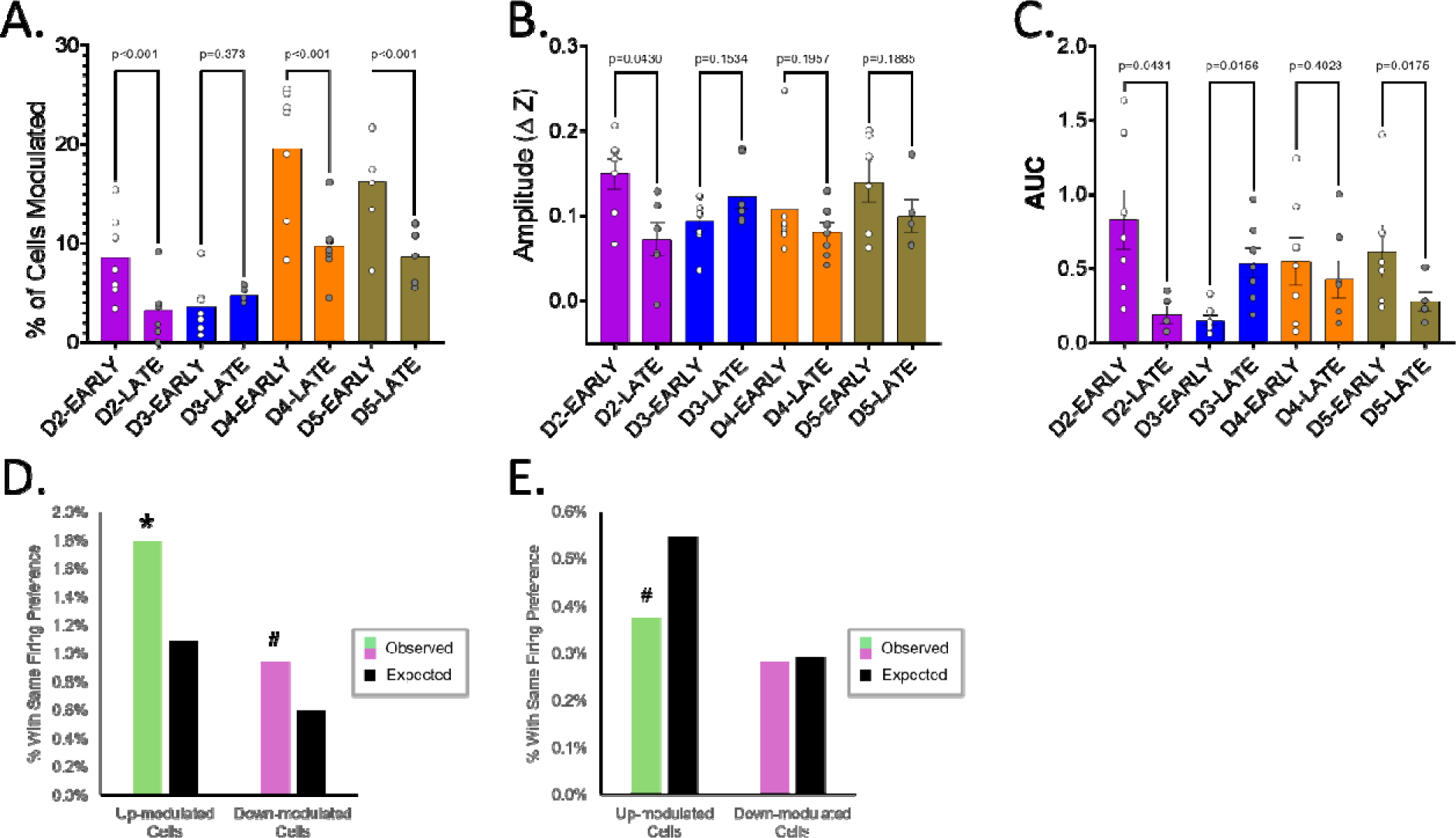
**A-C:** Graphs show dynamic within-sessions changes from early to late tones in (A) proportions of modulated neurons, (B) peak amplitude, and (C) response magnitude. **D:** Observed vs. expected proportions of neurons with the same firing preferences between the early tones of 24-h and 48-h test sessions. **E:** Observed vs. expected proportions of neurons with the same firing preferences between the early and late 24-h test tones. * p<0.05; # p<0.1.

We then tested recall-associated responses by comparing familiar but non-predictive tones from early acquisition to familiar but US-predictive tones from early 24-h test (a proxy for recall). EAPA response magnitude was greater during early 24-h test, along with a higher proportion of both up- and down-modulated neurons (LR cells; magnitude: p=0.0429; proportions: χ²=72.24, p<0.001; Figs. 4I-L). Thus, fear recall evoked a neural response driven by broader recruitment of ACC neurons.

Between-session comparisons across the 24-h and 48-h test sessions revealed that although EAPA responses and neuronal recruitment during the CS fluctuated, responses were not extinguished. Within both sessions, early tones engaged a greater fraction of modulated neurons than late tones (all cells; 24-h test: χ²=42.62, p<0.001; 48-h test: χ²=49.32, p<0.001), along with a decline in response magnitude from early to late tones of the 48-h test (p=0.0175; Figs. 5A-C). However, direct comparisons between the 24-h and 48-h tests showed no significant between-sessions changes in amplitude, magnitude, or proportions of modulated neurons (LR cells; amplitude: early-early p=0.45, late-late p=0.29; magnitude: early-early p=0.36, late-late p=0.41; proportions: early-early χ²=2.66, p=0.448, late-late χ²=1.85, p = 0.604; Figs. 4M-T). Thus, while ACC responses declined within sessions, consistent with a gradual reduction in CS representation in the absence of reinforcement, the reinstatement of the same early-late modulation profile across consecutive memory tests suggests a stable organizational pattern of cue encoding, concurrent with an absence of behavioral extinction.

In the pre-exposure and acquisition sessions, CS-responsive neurons transitioned symmetrically between modulation states, indicating a lack of individual neuron-level representation of the CS (McNemar’s tests, φ<0.093, p>0.05). However, in the 24-h test session, a specific subpopulation of CS-responsive neurons emerged that were up-modulated during the early 24-h test (i.e. recall) and preferentially retained their firing preferences in the early 48-h test (LR cells, N=1061, φ=0.097, p=0.039; Fig. 5D). These same cells trended toward a preferential transition to a non-modulated state from early to late 24-h test (LR cells, N=1061, φ=0.093, p=0.057; Fig. 5E) before being re-activated in the 48-h test session, a trend which was significant when analyzed using all within-session registered cells (all cells, N=2189, φ=0.094, p<0.001). This pattern of asymmetric transitions suggests that a specific subpopulation of up-modulated neurons represents the CS-US association during phases of recall when neural responses to the CS peak, and may be suppressed following neutral CS presentations.

### Changes in global and CS-evoked neural synchrony across sessions

To assess how ACC neural synchrony evolved with learning, we compared maximum pairwise correlations of CS event-aligned neural activity among longitudinally registered ACC neurons against a reference “Other” distribution of unaligned activity for each session (Fig. 6). For the cue pre-exposure and acquisition sessions (N=6 mice; Figs. 6A-C), the probability distribution functions of maximum pairwise correlations during the early and late tones showed no differences relative to the Other distributions (all p>0.1; Fig. 6B), indicating no overall difference in neural synchrony during CS presentations relative to unaligned activity. Nonetheless, there was a difference in neural synchrony between epochs (ANOVA, day*epoch*correlation interaction, F(19,95)=1.926, p=0.020), with the greatest neural synchrony during early cue pre-exposure. These differences were not specific to positive or negative correlations (Fig. 6C).

**Figure 6.**
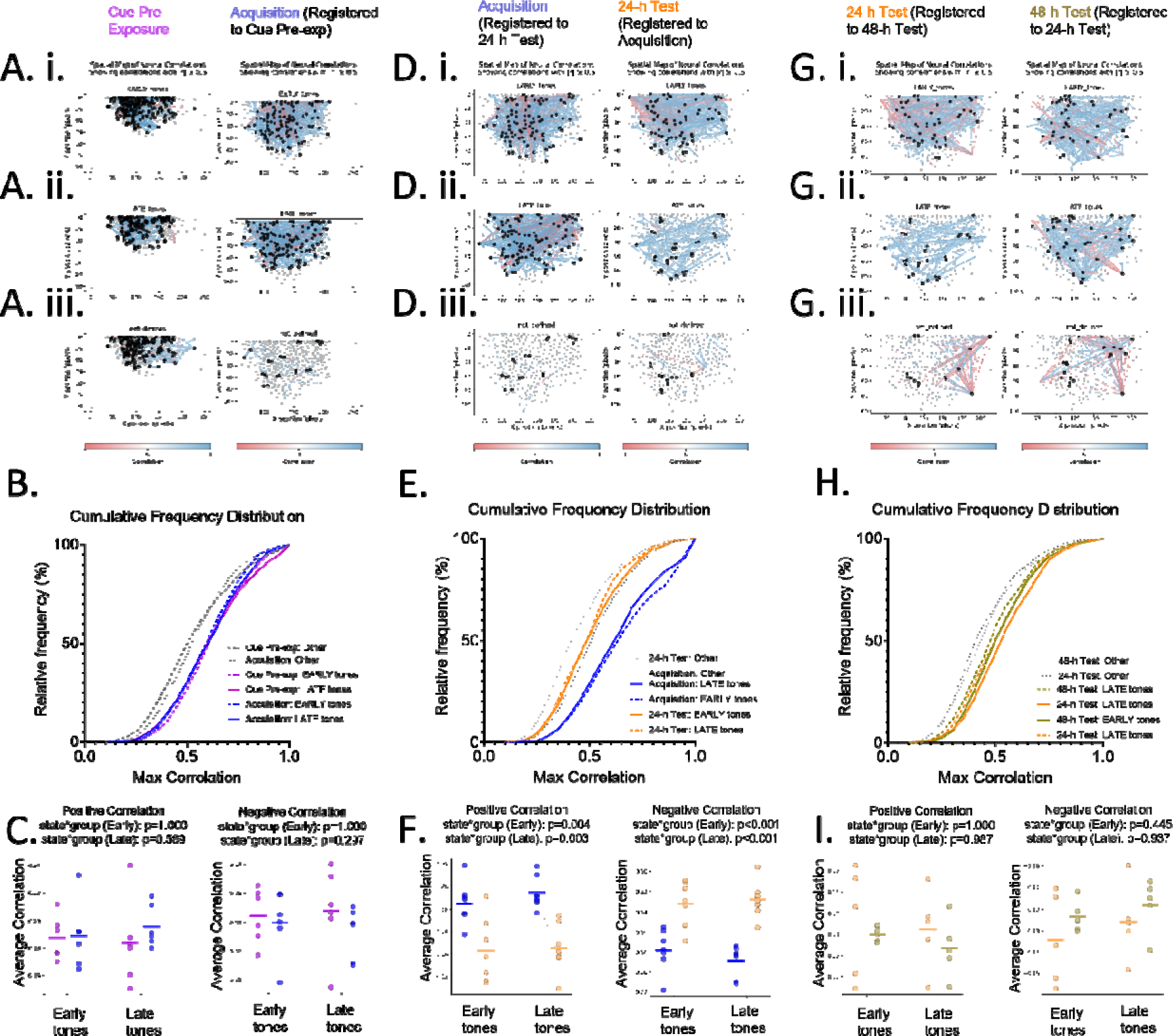
Global and CS-evoked neuronal synchrony during cue pre-exposure, acquisition, and memory tests. **A, D, G:** Spatial maps of neuron-to-neuron correlations for longitudinally registered cell pairs during (A) cue pre-exposure and acquisition, (D) acquisition and 24-h test, and (G) 24-h and 48-h test sessions. Maps show (i) early tones, (ii) late tones, and (iii) all tones combined. **B, E, H:** Cumulative frequency distributions (CFD) of maximum correlation values for early and late tones across (B) cue pre-exposure and acquisition, (E) acquisition and 24-h test, and (H) 24-h and 48-h test sessions, compared with unaligned “Other” activity. **C, F, I:** Comparisons of the positive (left) and negative (right) pairwise correlation values between early and late tones across (C) cue pre-exposure and acquisition, (F) acquisition and 24-h test, and (I) 24-h and 48-h test sessions.

We next compared neural synchrony during the acquisition and 24-h test sessions (N=7 mice; Figs. 6D-F). There was a significant change in neural synchrony during CS presentations between sessions (ANOVA, day*correlation interaction, F(19,114)=5.697, p<0.001). However, this effect coincided with a primary difference in the Other distributions between the two sessions (ANOVA, day*correlation interaction, F(19,114)=6.733, p<0.001), with a significant global decrease in neural synchrony in the 24-h test session. None of the individual epochs differed from the Other distributions with respect to neural synchrony (all p>0.1; Fig. 6E), indicating no differences between CS presentations and unaligned activity. This pattern was reflected in both the positive and negative correlations between sessions (Fig. 6F). These observations support the conclusion that consolidation was accompanied by a global reduction in ACC correlated activity and a weakening of ensemble coupling.

Finally, we examined whether synchrony continued to evolve between the 24-h and 48-h memory sessions (N=5 mice; Figs. 6G-I). Neural synchrony remained low and did not change throughout the 24-h and 48-h test sessions (ANOVAs, no interaction effects), either with respect to CS presentations or the Other distributions (Fig. 6H-I). These results indicate that the steep decline in synchrony occurred primarily during the consolidation interval and was not further changed between consecutive memory retrieval sessions.

Collectively, these results show that ACC synchrony peaked during early cue pre-exposure, then dropped dramatically and globally following consolidation, remaining globally reduced throughout the 24-h and 48-h test sessions.

## Discussion

This study was motivated by a knowledge gap regarding the specific role played by ACC in recall of a recent cued fear memory. We hypothesized that the ACC plays an active role in representing the negative valence of recently conditioned cues. By longitudinally recording ACC activity during fear learning and recall, we found that the ACC is indeed engaged during fear acquisition and acute fear recall in a manner consistent with encoding the negative valence of a CS. This encoding was dynamic and followed the predictive value of the CS, with neural responses increasing during acquisition, peaking during 24-h recall, and diminishing within sessions following repeated neutral CS presentations. Our interpretation is that the ACC encodes the dynamic change in valence of the CS, and may transmit this information to other regions containing the fear engram to enable dynamic updating. This interpretation can explain why the ACC and its projections to the engram-containing amygdala and hippocampus are necessary for cued learning processes, including acquisition (Bissière et al., 2008; Tang et al., 2005) and extinction (Fullana et al. 2018), but not for recent recall of cued fear (Frankland et al., 2004; Goshen et al., 2011), while helping to reconcile reports of ACC contributions to recent memory processes from other groups (Einarsson and Nader 2012; Bian et al. 2019; de Lima et al., 2022). However, future causal studies will be required to determine whether these patterns of changes in ACC neural responses are necessary for learning or behavior.

Our data do not indicate that the ACC supports recent fear recall through a robust or behaviorally dominant engram. Instead, we identified a sparse population of longitudinally registered ACC neurons that emerged during the 24-h memory test, selectively responded to early CS presentations, and maintained their firing preferences into the 48-h test. Although this population comprised only a small fraction of recorded neurons, its persistence across retrieval sessions suggests that these neurons may represent an early cortical trace associated with fear memory, rather than a fully established engram. Consistent with systems-level models of memory consolidation, cortical representations are thought to emerge gradually and initially remain behaviorally dispensable (Bontempi et al., 1999; Frankland & Bontempi, 2005, Kitamura et al., 2017). During this stage, these immature engrams coexist with hippocampal and amygdalar memory circuits before becoming functionally dominant at remote time points. Within this conceptual framework, the sparse ACC ensemble observed here may represent a nascent cortical trace that precedes the emergence of a fully developed ACC engram at remote time points, providing a cellular snapshot of early cortical engagement during systems consolidation.

Our analyses revealed that ACC population activity during freezing behavior is not homogeneous but instead organized into stable, intermingled subpopulations with opposing relationships to behavioral state. During both acquisition and 24-h recall, individual ACC neurons preferentially increased or decreased their activity during freezing relative to movement, forming distinct “Freezing On” and “Freezing Off” subpopulations. The state preferences of these subpopulations were expressed at levels exceeding chance and were maintained across sessions, indicating that they reflect stable functional identities rather than transient fluctuations. At the population level, freezing during CS presentations was associated with reduced mean ACC activity and increased neural synchrony, suggesting a coordinated reconfiguration of network dynamics during fear expression that favors transmission of a learning signal through increased signal-to-noise ratio. This is consistent with other studies that have identified ON and OFF neuronal ensembles in prefrontal cortex transmitting information about novelty and salience during social interactions (Liang et al., 2018).

Behaviorally, freezing was elevated during “Late” tones of acquisition and “Early” tones during the 24-h test, and remained stable over 48-h. While the freezing state was associated with decreased neural activity and increased neural synchrony, CS responses peaked during early 24-h and 48-h epochs, and the 24-h and 48-h test sessions were associated with a global decrease in neural synchrony. Therefore, responses to the CS cannot be driven by a greater proportion of freezing, or vice versa. Freezing onset events, whether occurring during CS presentation or during tone-free periods, did not evoke consistent ACC responses, and the proportions of modulated neurons were indistinguishable across contexts. Therefore, responses to the CS cannot be driven by the relative proportion of freezing (or vice versa) or by the cessation of movement. suggesting that responses were not driven by cessation of movement inherent to freezing, or to the relative proportion of freezing. These results demonstrate that ACC modulation during acquisition and recall is tied specifically to the associative value of the CS, rather than to freezing behavior, motor movement, or novelty.

We also found that the ACC encodes for novelty of both the CS and the US, consistent with the known role of the ACC in novelty and prediction error (Alexander & Brown, 2019; Brown & Braver, 2005; Hayden et al., 2011; Ito et al., 2003; Sarafyazd & Jazayeri, 2019; Wessel et al., 2014). CS responses peaked during initial presentations in early cue pre-exposure, then rapidly attenuated with subsequent, neutral presentations. Similarly, neural responses to the initial US were higher than subsequent US presentations with respect to response amplitude and proportions of up-modulated neurons. Indeed, within our own results, averaging across entire sessions instead of by “early” and “late” epochs showed no differences in neural response to the CS between cue pre-exposure and acquisition, due to the confounding representation of novelty in one session and valence in the other. These findings may explain, in part, previous observations of weak differences in neuronal response across CS presentations in cued fear acquisition paradigms without cue pre-exposure (Steenland et al., 2012). The fear conditioning paradigm serves as the cornerstone for studying the learned association between cues and threatening events (Fendt & Fanselow, 1999; Kim & Jung, 2006; Maren, 2001), and these results emphasize the need to control for novelty in experiments intended to look for learning effects in the ACC.

The findings in this study should be considered in light of its limitations. First and foremost, as a fully within-subjects study, this study is incapable of establishing cause-effect relationships among the observed effects. While we have shown a pattern of responses in the ACC consistent with the dynamic encoding of negative valence, we have not manipulated those responses to demonstrate that they affect behavioral outcomes, although prior literature supports that conclusion (Bissière et al., 2008; Tang et al., 2005). Because all female subjects were eliminated from our cohort, our study includes only male subjects, and may not fully describe these same representations in females. We did not manipulate the context in our design, so the memory sessions represent overlapping contextual and cued fear responses. Our “early” and “late” time points during cue pre-exposure and acquisition contained only two tones each, increasing the random variability in responses. Finally, the absence of behavioral extinction despite repeated unreinforced presentations of the CS in our study may reflect insufficient tone numbers or the use of a consistent context. Although we reported the percentage of significantly modulated neurons as one index of cue-evoked activity, we note that this measure can be misleading when taken in isolation. For example, in some conditions the fraction of modulated neurons was close to 5%, yet the averaged population trace was essentially flat, while in other conditions a comparable fraction coincided with a clear population waveform. This discrepancy likely reflects that population amplitude depends not only on the number of cells passing threshold but also on their magnitude and temporal alignment. Synchrony and population-level amplitude therefore likely provide more sensitive indicators of network engagement than the proportion of individually significant neurons, which is strongly shaped by statistical thresholding.

In conclusion, this study highlights the dynamic role of the ACC in detecting novelty, representing aversive outcomes, and representing the negative valence of a conditioned cue as it changes dynamically across cued fear acquisition, consolidation, recall, and extinction. By dissociating novelty from associative value, we identified distinct patterns of ACC modulation that reflect both prediction error signals and network plasticity. We identified stable populations of “freezing on” and “freezing off” neurons in the ACC that participated in increased neural synchrony during freezing. Additionally, we identified a sparse and stable subpopulation of ACC neurons that emerged during early retrieval that may represent an incipient cortical memory trace, consistent with models of gradual systems consolidation in which cortical engrams mature over time. Our findings are consistent with theoretical models positioning the ACC as a site for novelty, aversion, and outcome evaluation, and underscore its role in continuously updating threat appraisal by dynamically representing the changing valence of a cue.

## Acknowledgements

We gratefully acknowledge Shay Neufeld, Senior Director of Data Products and Analytics at Inscopix, for implementing several modifications to the IDEAS platform to facilitate the analysis of data presented in this study. We would like to thank Dr. Sarah Daniel of In Scripto (inscriptoscience.com) for scientific writing consultation on this manuscript.

## Author Contributions

PG and JPB designed the overall research study, including the conceptual background and experimental design. PG performed all GRIN lens implant surgeries, behavioral testing and data acquisition. NS assisted with behavioral setup during data acquisition and performed analysis of animal behavior. RS performed histological verification and fluorescence imaging. JSS contributed to animal surgery and behavioral procedures under PG’s supervision. KG contributed to data quality control and preprocessing under PG’s supervision. KLN provided experimental, administrative, and managerial support. RAP advised on calcium imaging analysis and early data processing strategies. PG and JPB performed all calcium imaging data analysis and prepared all figures. PG, RAP, and JPB collectively interpreted all results. PG prepared the initial draft of the manuscript, and PG, RAP, and JPB revised the manuscript into final form. All authors reviewed and approved the final manuscript.

## Data and Materials Availability

All raw data and metadata are available on Figshare (doi: TBD). All custom data analysis scripts are available on Github (doi: TBD).

## Supplementary Materials

Supplementary figures and tables can be found in the Supplemental Materials file.

---

**Supplementary Figure 1.**
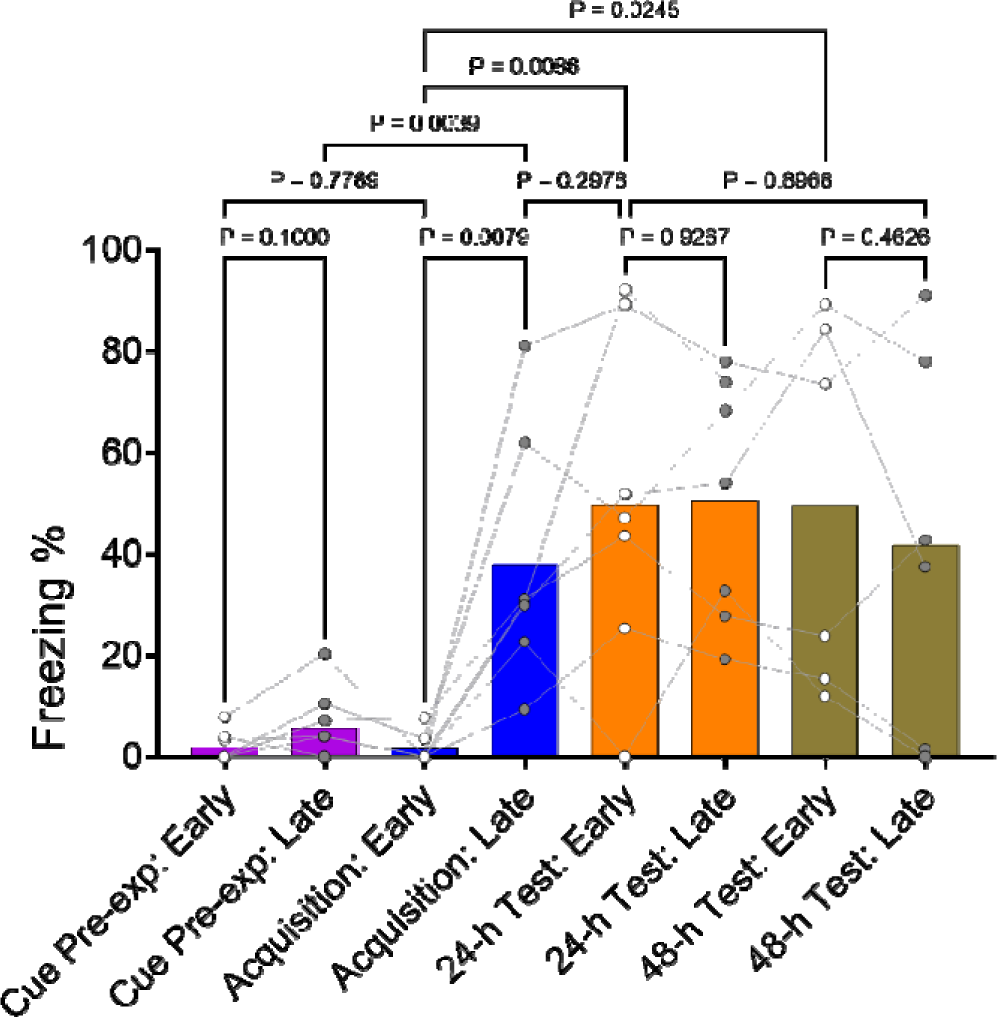
Freezing behavior across epochs of the cued fear paradigm. Quantification of freezing behavior across the defined early and late tone epochs for each session, shown as mean ± SEM with individual subject data overlaid.

**Supplementary Figure 2.**
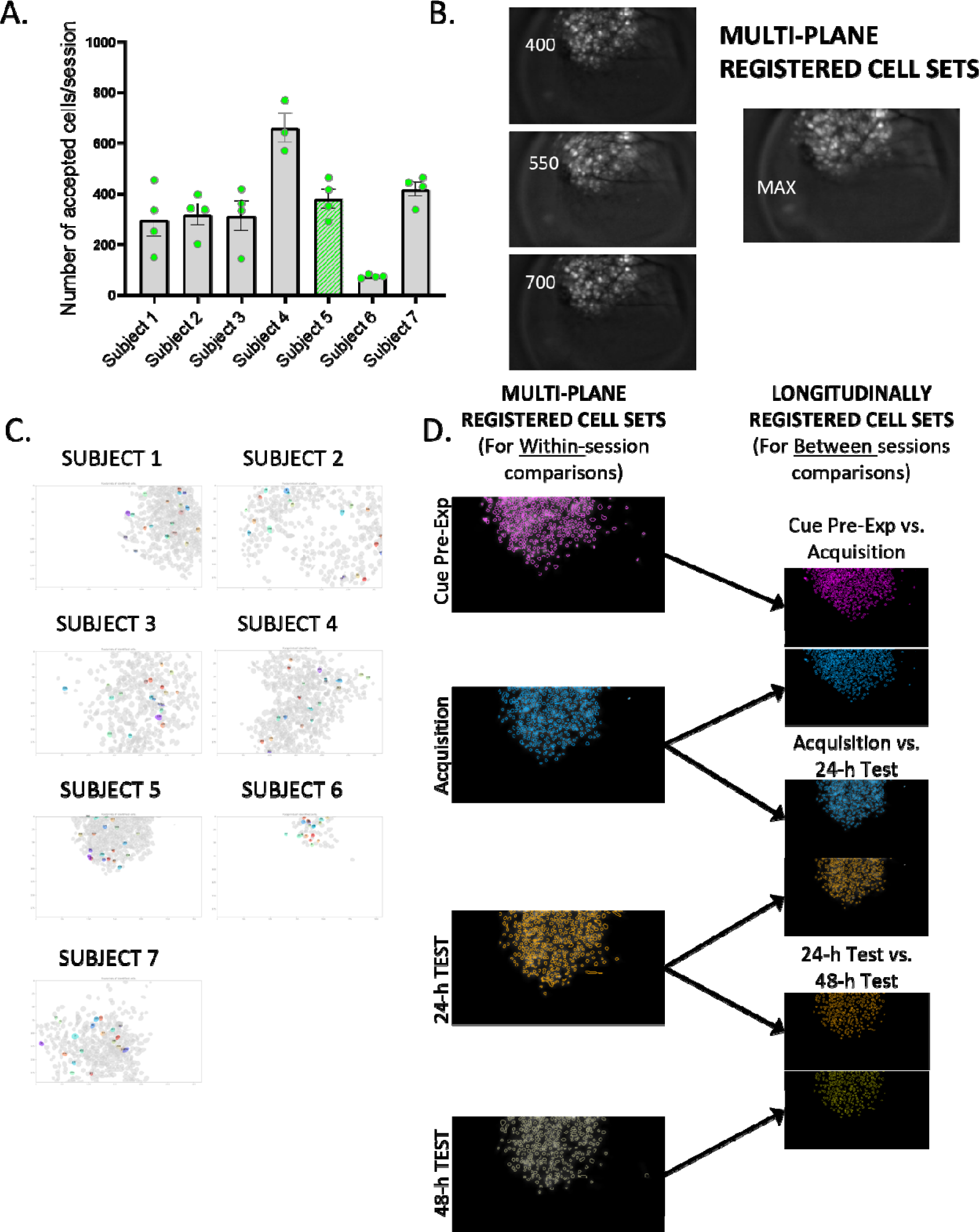
Calcium imaging and registration strategy in mouse ACC. **A:** Number of accepted neurons per session across the seven animals included in the study (mean ± SEM: ~300–400 neurons/session per subject). The green-highlighted bar indicates the representative subject shown in the main figures throughout the manuscript. **B:** Multi-plane registration across three focal depths (400, 550, and 700 units) was used to align spatial footprints and generate a unified cell set for each subject. The composite image (MAX) shows the merged footprint map across planes. **C:** Spatial footprints of all accepted neurons during the cue pre-exposure session for each individual subject. **D:** Schematic of analysis strategy for within- and between-session comparisons. Left: within-session analyses included all multi-plane–registered neurons from a given session. Right: between-session analyses used only longitudinally registered neurons tracked across two consecutive sessions.

**Supplementary Figure 3.**
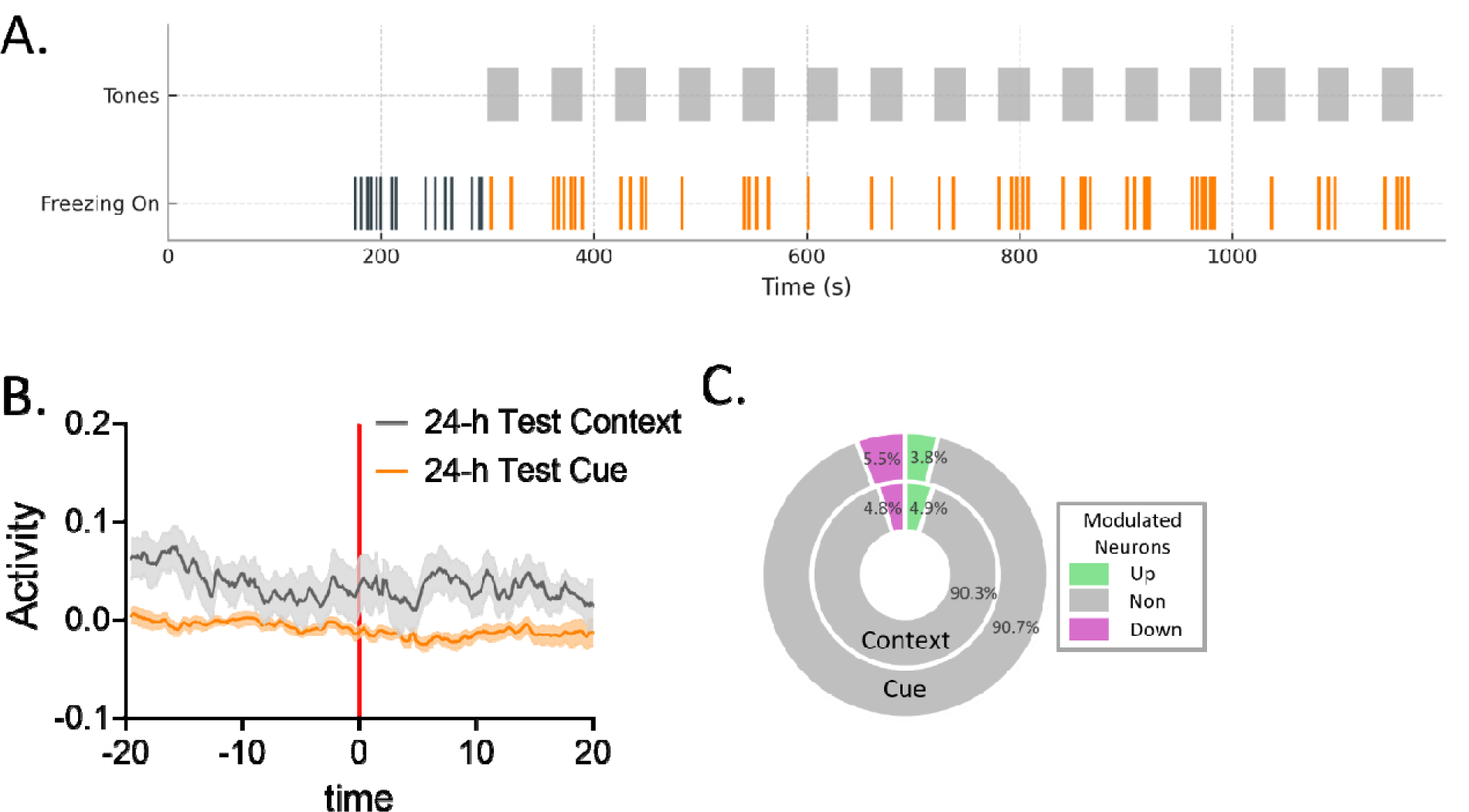
ACC activity at freezing onsets during contextual and cued freezing event conditions in the 24-h test. **A:** Representative raster plot showing the timing of tone presentations (gray) and freezing onsets (orange) during the 24-h test session. **B:** Averaged peri-event population activity aligned to freezing onset, comparing freezing bouts occurring in the absence of tones (context, black) versus those during tone presentation (cue, orange). Red line marks freezing onset (t = 0). **C:** Proportions of up-modulated (green), down-modulated (purple), and non-modulated (gray) neurons during context-related freezing onsets and cue-related freezing onsets, revealing no significant differences in recruitment across conditions.

**Supplementary Figure 4.**
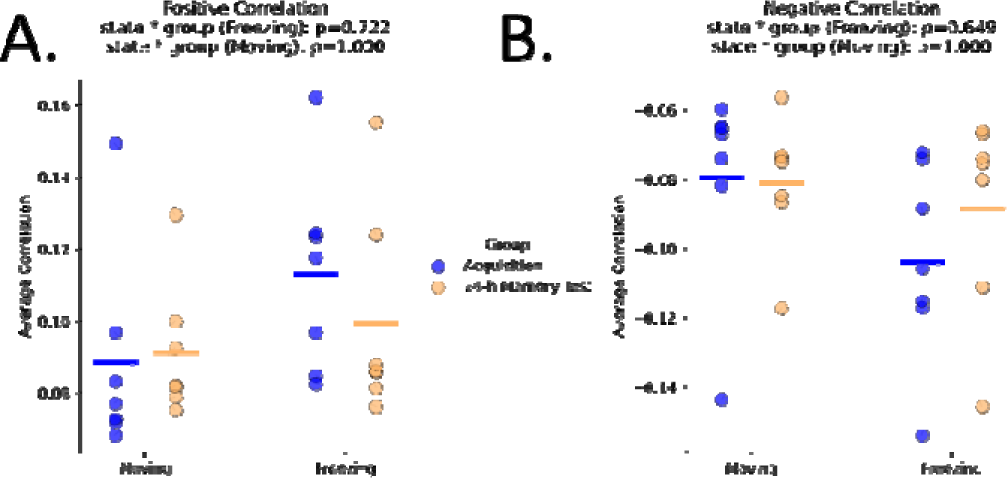
**A-B:** Comparisons of the (A) positive and (B) negative pairwise correlation values between freezing and moving states in the acquisition and 24-h test sessions.

**Supplementary Figure 5.**
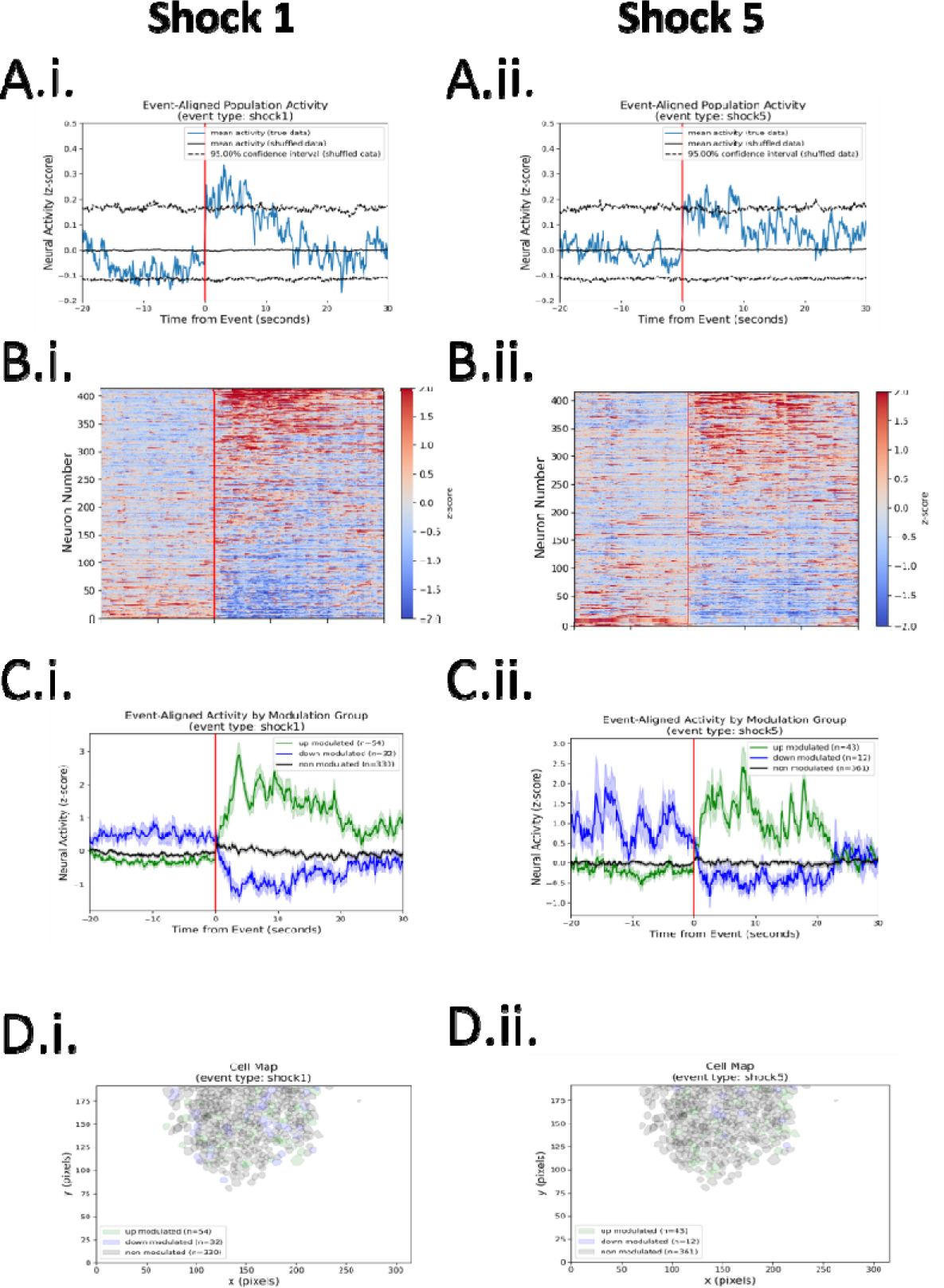
Comparison of ACC neural activity in response to the first and last US from a representative subject. **A:** Event-aligned population activity traces showing change of population activity over time in response to the (i) first and (ii) fifth US events during the acquisition session. Red line represents timing of the delivery of the US events (t=0). **B:** Heatmaps of event-aligned neural activity in response to the (i) first and (ii) fifth US events during acquisition, each row representing individual neuron responses aligned to the time of the event. **C:** Event-aligned activity traces for up-modulated (green), down-modulated (blue), and non-modulated (gray) neurons during the (i) first and (ii) fifth US events. **D:** Cell maps representing the spatial distribution of neurons modulated by the (i) first and (ii) fifth US events.

**Supplementary Figure 6.**
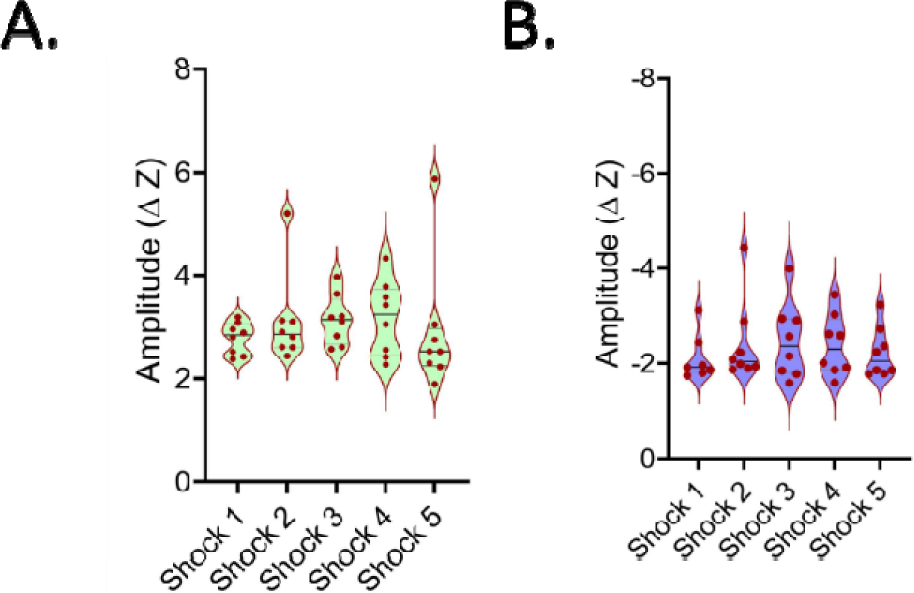
Violin plots of peak response amplitudes for (A) up-modulated and (B) down-modulated neurons across US events. All error bars are SEM.

**Supplementary Figure 7.**
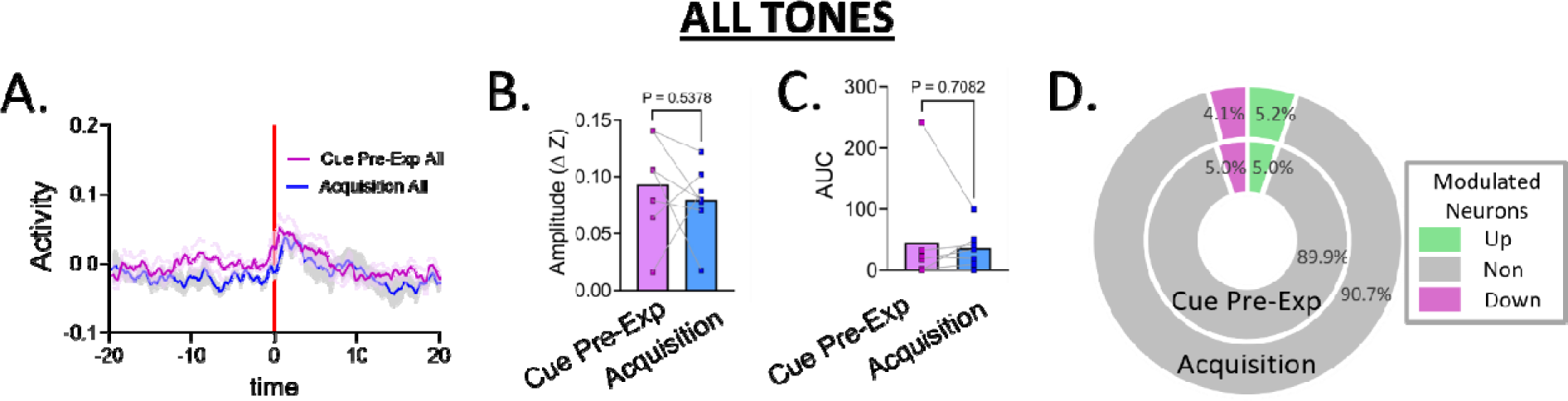
ACC responses to all tones during cue pre-exposure and acquisition. **A:** EAPA traces for all tone presentations during cue pre-exposure (magenta) and acquisition (blue). Red line marks tone onset (t = 0). **B-C:** Quantification of (B) peak amplitude and (C) response magnitude revealed no significant differences between cue pre-exposure and acquisition (all p > 0.5). **D:** Proportions of up-modulated (green), down-modulated (purple), and non-modulated (gray) neurons during cue pre-exposure and acquisition, showing no differences in modulation.

